# Distinct signalling routes mediates intercellular and intracellular rhizobial infection in *Lotus japonicus*

**DOI:** 10.1101/2020.05.29.124313

**Authors:** Jesús Montiel, Dugald Reid, Thomas H. Grønbæk, Caroline M. Benfeldt, Euan K. James, Thomas Ott, Franck A. Ditengou, Marcin Nadzieja, Simon Kelly, Jens Stougaard

**Author notes:** Correspondence: Simon Kelly and Jens Stougaard. **Contributions**, J.M., D.R., C.M.B. and T.H.G. characterised and isolated the mutants. D.R. and J.M. processed and analysed the RNAseq data, respectively. M.N. and E.K.J. performed the microscopy analyses. J.M. wrote the manuscript and prepared the figures with contributions of the co-authors. J.M., S.K. and J.S. conceived the research plan. J.S. and S. K. coordinated and guided the research.

## Abstract

Rhizobial infection of legume roots during development of nitrogen fixing root nodules occurs either intracellularly though plant derived infection threads traversing the epidermal and cortical cell layers to deliver the bacteria or intercellularly via bacterial entry between epidermal plant cells. Although, around 25% of all legume genera are postulated to be intercellularly infected, the pathways and mechanisms supporting this process has remained virtually unexplored due to lack of genetically amenable legumes that have this infection mode. In this study, we report that the model legume *Lotus japonicus* is infected intercellularly by *Rhizobium* sp. IRBG74 and demonstrate that the resources available in *Lotus* enable insight into the genetic requirements and the fine-tuning of the pathway governing intercellular infection. Inoculation of *Lotus* mutants shows that *Ern1* and *RinRK1* are dispensable for intercellular infection in contrast to intracellular infection. Other symbiotic genes, including *Nfr5, SymRK, CCaMK, Epr3, Cyclops, Nin, Nsp1, Nsp2, Cbs* and *Vpy1* are equally important for both entry modes. Comparative RNAseq analysis of roots inoculated with IRBG74 revealed a distinctive transcriptome response compared to intracellular colonization. In particular, a number of cytokinin-related genes were differentially regulated. Corroborating this observation *cyp735A* and *ipt4* cytokinin biosynthesis mutants were significantly affected in their nodulation with IRBG74 while *lhk1* cytokinin receptor mutants did not form any nodules. These results indicate that a differential requirement for cytokinin signalling conditions intercellular rhizobial entry and highlight the distinct modalities of the inter- and intra-cellular infection mechanisms.

## Introduction

Legumes constitute a large and diverse plant family and most legumes are able to develop nitrogen fixing root nodules in symbiosis with soil bacteria commonly referred to as rhizobia. Bacterial infection of roots and root nodules through intracellular infection threads has been extensively researched in the model legumes *Lotus japonicus* (*Lotus*) and *Medicago truncatula* (*Medicago*) as well as crop legumes like soybean (*Glycine max* L.), pea (*Pisum sativum* L.) and common bean (*Phaseolus vulgaris* L.). However, an alternative mechanism of intercellular infection is widespread in different genera of the Fabaceae family (Sprent et al., 2017) and suggested to be an ancient and less sophisticated mechanism of rhizobial colonization (Sprent, 2007). In this process, rhizobia invade the legume roots between epidermal/root hair cells or by crack entry, during the protrusion of lateral roots (Coba de la Pena et al., 2017). Intercellular infection processes has been described in detail by microscopy in different legumes such as *Mimosa, Neptunia, Stylosanthes, Cytisus* and *Lupinus* (de Faria et al., 1988; James et al., 1992; Subba-Rao et al., 1995; Vega-Hernandez et al., 2001; Gonzalez-Sama et al., 2004; Goormachtig et al., 2004). Special attention has been dedicated to characterise the histology of intercellular infection and nodulation processes in the temperate legumes *Arachis hypogaea, Aeschynomene* sp. and the semiaquatic legume *Sesbania rostrate* (Chandler, 1978; Boogerd and Van Rossum, 1997; Bonaldi et al., 2011; Ibanez et al., 2017). Under flooded conditions, rhizobial colonization takes places via infection pockets in *S. rostrata*, formed by a cell death process that depends on nodulation factors (NFs), perception and localized formation of reactive oxygen species (D’Haeze et al., 2003). From such infection pockets, cortical infection threads are formed and migrate to the nodule primordium, where the bacteria are released from the infection threads (ITs) and colonize the nodule cells in symbiosomes (Capoen et al., 2010). Unlike intracellular colonization, the leucine-rich repeat-type receptor kinase *SymRK* gene and the Ca2+/calmodulin-dependent protein kinase *CCaMK* gene are dispensable for intercellular infection in *S. rostrata* by *Azorhizobium caulinodans*. However, these genes are required for the subsequent intracellular infection threads (Capoen et al., 2005; Capoen et al., 2009). In peanut (*A. hypogaea*), Bradyrhizobia enter through the middle lamellae of two adjacent root hairs and spread intercellularly between epidermal and cortical cells. In parallel, adjacent axillary root hair basal cells become enlarged and infected by the microsymbiont (Chandler, 1978; Boogerd and Van Rossum, 1997; Guha et al., 2016). Both nodule formation and nodule cell colonization require proper exopolysaccharide production by rhizobia (Morgante et al., 2007). However, invasion and nodule organogenesis can be achieved in peanut by rhizobia lacking *nod* genes (Ibanez and Fabra, 2011; Guha et al., 2016). Similarly, a NF-independent nodulation program has been described in certain *Aeschynomene* spp. (Giraud et al., 2007). Currently, the *A. evenia-Bradyrhizobium* symbiosis has been employed to study the molecular genetics of this unusual NF-independent symbiosis (Arrighi et al., 2012). Recent findings show that during this peculiar mechanism, several components of the NF-dependent process are also recruited, such as SYMRK, CCaMK and the histidine kinase HK1 cytokinin receptor (Fabre et al., 2015). Additionally, the structural requirements to perceive NFs in *S. rostrata* are more permissive in intercellular infection compared to the intracellular infection (Goormachtig et al., 2004). Intercellular infection occurs in several *Sesbania* spp. by IRBG74 (Cummings et al., 2009), a nodulating *Rhizobium* sp. strain that is also able to colonize rice and Arabidopsis roots as an endophyte (Biswas et al., 2000; Mitra et al., 2016; Zhao et al., 2017). Therefore, a better understanding of intercellular colonization would facilitate the engineering of non-legume crops for colonization by nitrogen-fixing bacteria.

The better characterised intracellular infection is dependent on rhizobial signal molecules, lipochitooligosaccharides, called nodulation factors (NFs). In *Lotus*, the NFs are recognized in the plasma membrane of the root hairs by Nod Factor receptors (NFR1, NFR5 and NFRe; (Madsen et al., 2003; Radutoiu et al., 2003; Murakami et al., 2018). A compatible recognition leads to rhizobial attachment to the root hair tip, promoting its curling to trap the bacteria with an infection pocket. This give rises to the formation of an infection thread (IT), a tubular structure with an inward growth that originates from invagination of the plasma membrane of the root hair. The IT follows a polar growth towards inner root cell layers, reaching the nodule primordia, formed by the activation of cell division in the cortical cells. The nodule primordia give rise to a mature nodule, where the bacteria are released from the IT into symbiosomes and differentiate to become nitrogen-fixing bacteroids (Downie, 2014).

The intracellular infection of rhizobia *via* ITs in the root hairs has been extensively investigated in *Lotus* and *M. truncatula* (Lace and Ott, 2018). In these legumes, the infection is orchestrated by several transcription factors, including Nodule Inception (*Nin*; (Schauser et al., 1999; Marsh et al., 2007) *Nsp1/Nsp2* (Kalo et al., 2005; Heckmann et al., 2006), *Cyclops* (Yano et al., 2008; Singh et al., 2014) and *ERF* Required for Nodulation (*Ern1*; (Cerri et al., 2012; Kawaharada et al., 2017). The latter is required for activation of the expression of the cytokinin-biosynthesis genes *Ipt2* and *Log4*, that are major contributors to the initial cytokinin symbiotic response in *Lotus* (Reid et al., 2017). Cytokinin is necessary for nodule organogenesis, but plays a negative role during rhizobial invasion. In the cytokinin oxidase/dehydrogenase 3 mutant (*ckx3*), where cytokinin levels in the roots are increased, rhizobial infection is significantly reduced (Reid et al., 2016). By contrast, the roots of cytokinin receptor *Lhk1* mutants are hyperinfected by rhizobia (Murray et al., 2007). Recent reports show that other receptors have a positive role in infection, like the exopolysaccharide receptor EPR3 (Kawaharada et al., 2015) and the Leu-rich repeat receptor-like kinase RINRK1 (Li et al., 2019). In addition, several molecular components are required. Mutants disrupted in the E3 ligase *Cerberus* (Yano et al., 2009), the nodule pectate lyase *Npl1* (*Xie et al., 2012*) or *Arpc1, ScarN, Nap1, Pir1* involved in actin rearrangements (Yokota et al., 2009; Hossain et al., 2012; Qiu et al., 2015), show defects in IT development and abortion of the infection process. In *Medicago* the infection thread localized Cystathionine-β-Synthase-like 1 (CBS1; (Sinharoy et al., 2013)), coiled-coil RPG protein (Arrighi et al., 2008) and Vapyrin (Murray et al., 2011) are crucial components of the root hair infectome (Liu et al., 2019).

In a first step to compare the genetic programs controlling intracellular and intercellular infection we have explored the infective capacity of *Rhizobium* sp. strain IRBG74 in *Lotus*, where a wide range of genetic, genomic and transcriptomic resources are available. Crucial genes for both modalities of rhizobial infection were identified along with distinctive cellular, transcriptome and genetic requirements for the intercellular colonization.

## Results

### IRBG74 induces nitrogen-fixing nodules in *Lotus*

The rhizobial strain IRBG74, infects *S. cannabina* intercellularly (Cummings et al., 2009; Mitra et al., 2016) and interestingly it is also capable of colonizing *O. sativa* and *A. thaliana* roots as an endophyte (Mitra et al., 2016; Zhao et al., 2017). In order to evaluate the infective capacity of IRBG74 in *Lotus*, nodulation kinetics were recorded from 1 to 6 weeks post-inoculation (wpi) of the Gifu accession. As a control, *L. japonicus* seedlings were inoculated with its customary symbiont *Mesorhizobium loti* R7A, that infects intracellularly. Nodule primordia were observed at 1 wpi on *Lotus* roots inoculated with *M. loti*, with mature pink nodules evident by 2 wpi (Fig. 1A). IRBG74 also induced nodule organogenesis in *Lotus*, but the first nodule primordia were not observed until 2 wpi (Fig. 1A and B). The first mature pink nodules on plants inoculated with IRBG74 usually appeared at 3 wpi, but these were evidently smaller compared to the pink nodules induced by *M. loti* at the same time point (Fig. 1B). In addition, the number of pink and white nodules were significantly lower at 2 and 3 wpi in plants inoculated with IRBG74 compared to plants inoculated with *M. loti* (Fig. 1A). After 4 to 6 wpi the number of nodules on plants inoculated with *M. loti* and IRBG74 was comparable (Fig. 1A and C). The delay in the IRBG74 nodulation was reflected by the distribution pattern of the nodules in the root system, since 45% of the nodules were found in the root crown (upper 2 cm of the root system), while plants inoculated with *M. loti* had 84% of the nodules in this segment of the root (Fig. 1C and D). Nodules induced by IRBG74 were pink and the plant shoots were green (Fig. 1B and C), indicative of nitrogen fixation. At 6 wpi, fresh weight and shoot length were significantly lower, compared to plants inoculated with *M. loti* reflecting the delayed nodule formation (Fig. 1E and F). These results show that IRBG74 is able to induce nitrogen-fixing nodules in *Lotus*, albeit with a delay.

**Figure 1.**
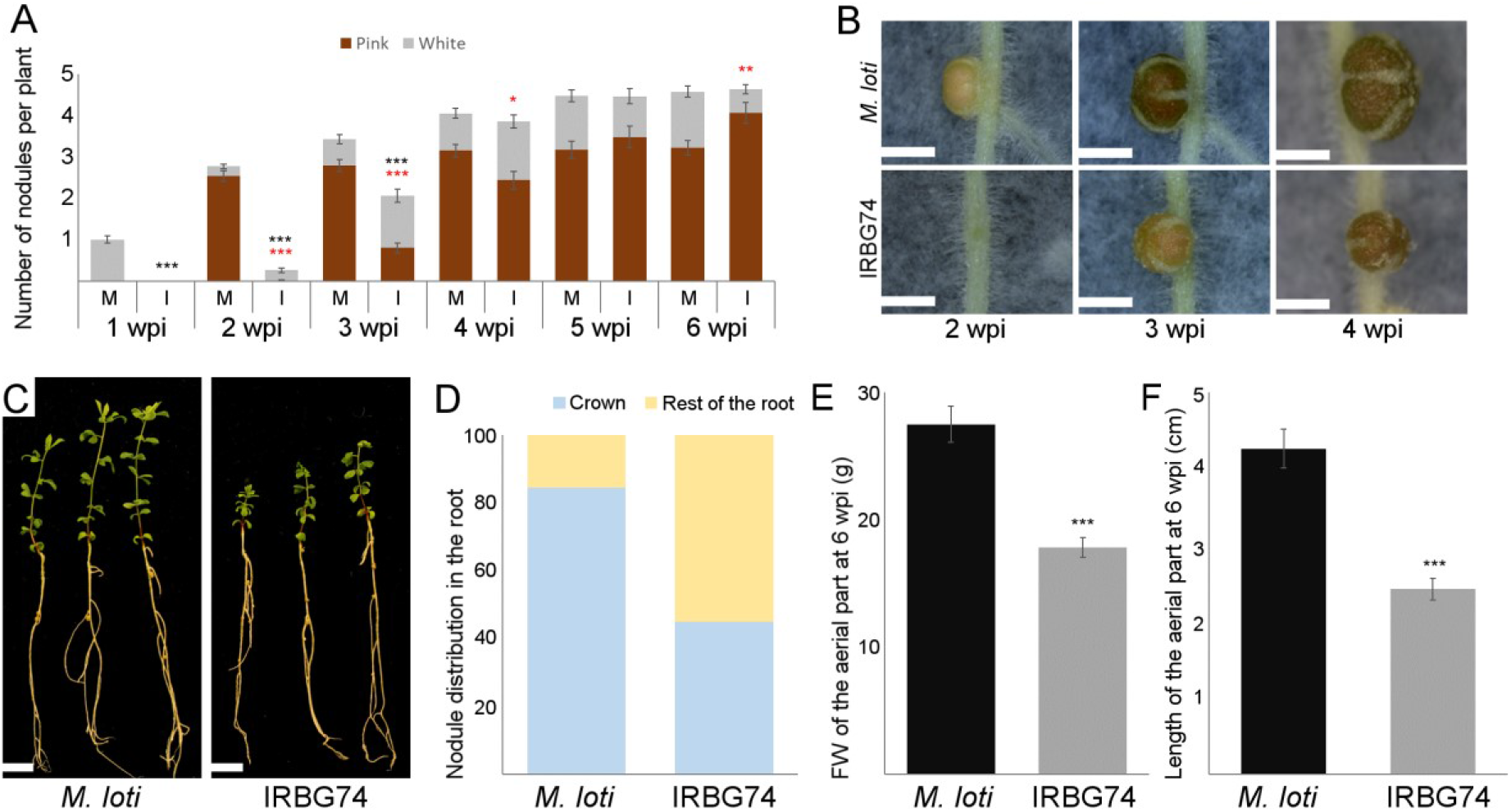
Nodulation phenotype of *Lotus* plants inoculated with *M. loti* (M) or IRBG74 (I). A, Nodule numbers at 1 to 6 wpi. B, Images of nodules at 2-4 wpi. C-F, Distribution of nodules in the root system, fresh weight and shoot length at 6 wpi. Scaler bar, 1 mm (B) and 1 cm (C). Student’s *t*-test of pink and total number of nodules (red and black asterisks, respectively), FW and length of the aerial part between plants inoculated with *M. loti* and IRBG74. P-values < 0.05, 0.01 and 0.001 are marked with one, two or three asterisks, respectively. n = 68 (M), 66 (I).

### *Lotus* is intercellularly infected by IRBG74

The delay in organogenesis following inoculation with IRBG74 compared to *M. loti* prompted us to explore the infection process. For this purpose, the constitutive DsRED expressing plasmid pSKDSRED was transformed into IRBG74 in order to monitor the early infection process by confocal microscopy. *M. loti*-DsRed was used as a control. Infection and nodule organogenesis were unaltered with these engineered strains. Typical intracellular ITs in long root hairs were abundant at 7 dpi with *M. loti*-DsRed (Fig. 2A and D). In contrast, no IRBG74-DsRed ITs were observed. IRBG74 was attached to the surface of the roots, mainly associated with the boundaries of the epidermal cells (Fig. 2B). At 2 wpi IRBG74-DsRed invaded cortical root cell layers intercellularly (Fig. 2C; Supplemental Movie S1). A detailed and quantitative inspection revealed an average of 35 root hair infection threads in response to *M. loti*-DsRed, while none were found in the roots inoculated with IRBG74-DsRed at 10 and 21 dpi (Fig. 2D).

**Figure 2.**
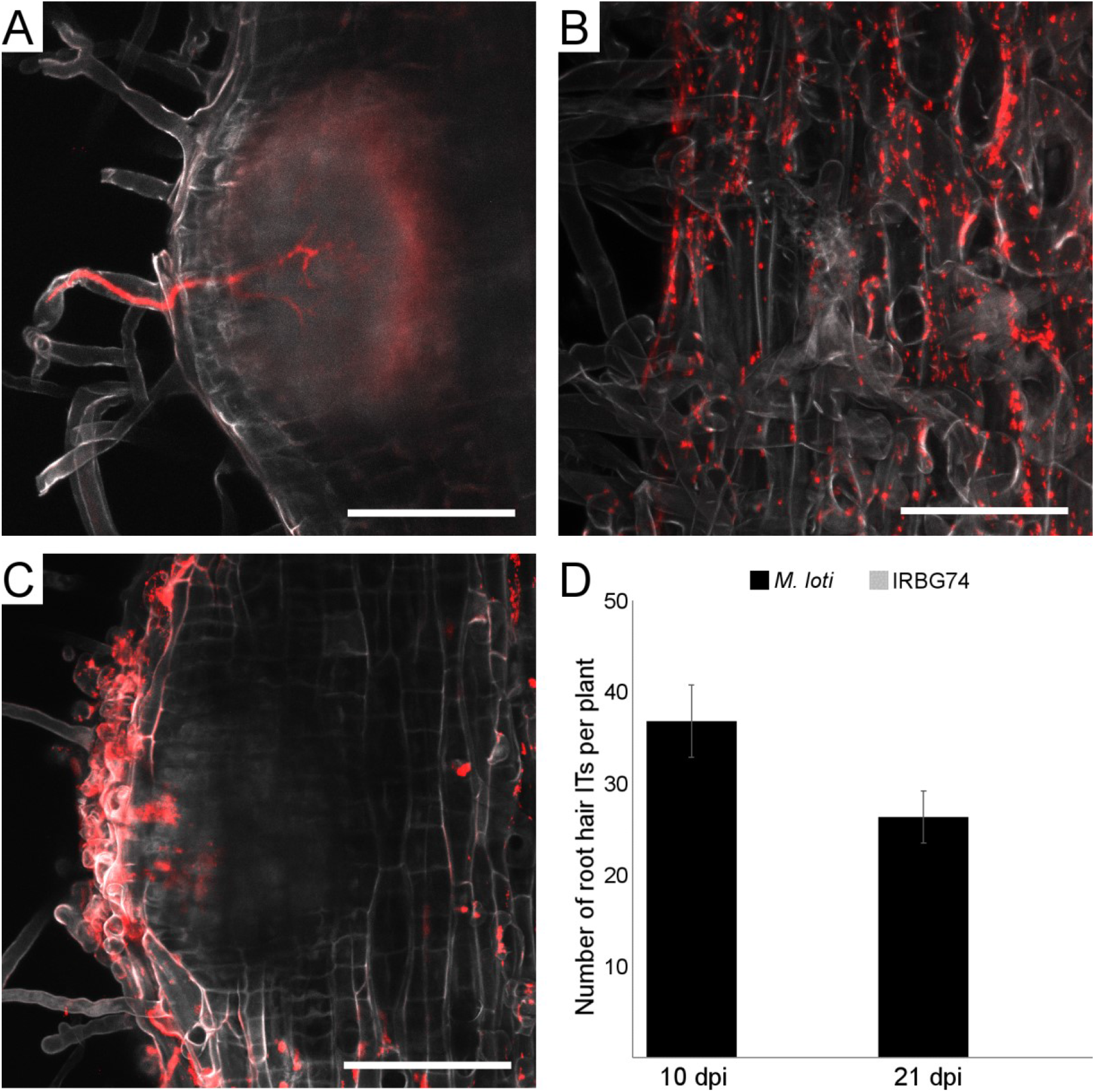
Intracellular and intercellular rhizobial infection of *Lotus* roots. Confocal microscopy images of *Lotus* roots at 1 wpi (A and B) and 2 wpi (C) with *M. loti*-DsRed (A) or IRBG74-DsRed (B and C). Scale bar, 50 μm. D, number of root hair ITs in *Lotus* plants at 10 dpi and 21 dpi with *M. loti*-DsRed or IRBG74-DsRed (n ≥ 15). Error bars mean SEM.

In order to characterize the progression of the IRBG74 infection process, the histology of young and mature nodules was analysed by light microscopy and compared to nodules of similar developmental stage induced by *M. loti* at 3 wpi. Symbiosome-containing nodule cells and transcellular ITs were observed in both young and mature nodules of *Lotus* with both rhizobial strains (Fig. 3A-D; Supplemental Fig. S1). The transcellular ITs were remarkably more numerous in nodules colonized by *M. loti*, compared to IRBG74, particularly in young nodules (Fig. 3E).

**Figure 3.**
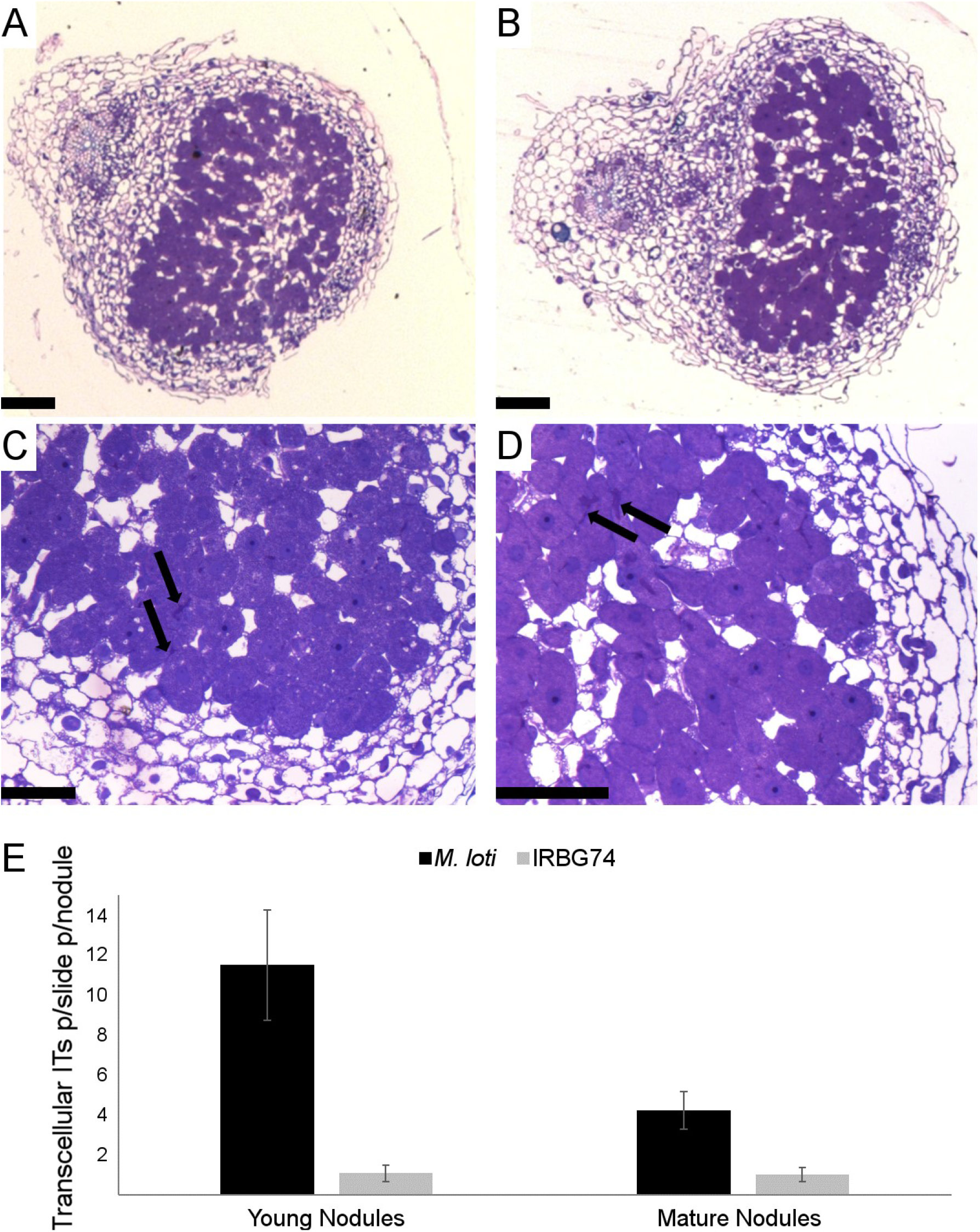
Histology of *Lotus* nodules colonized by *M. loti* or IRBG74. Sections from young nodules of *Lotus* inoculated with *M. loti* (A and C) or IRBG74 (B and D) were stained with tolouidine blue to evaluate rhizobial occupancy in the nodule cells. Transcellular infection threads are indicated with arrows. Scale bar, 100 μm (A and B) and 50 μm (C and D). E, number of transcellular ITs in 7 slides from 7 different young (Y) and mature (M) nodules. Error bars mean SEM.

### *rinrk1* and *ern1* mutants show contrasting symbiotic phenotypes with IRBG74 and *M. loti* inoculation

Root infection and nodule organogenesis are highly coordinated multistep processes and nodule organogenesis is affected in mutants interrupted in rhizobial infection (Oldroyd and Downie, 2008; Madsen et al., 2010). Nodulation kinetics was therefore a suitable readout to determine the genetic dependency of the *Lotus*-IRBG74 intercellular process. IRBG74 and *M. loti* induced nodulation of a set of previously identified *Lotus* symbiotic mutants was scored at 1-6 wpi. First, mutants affected in the symbiotic receptor genes *Nfr5, SymRK, RinRk1* and *Epr3* were tested. Both *M. loti* and IRBG74 were unable to form nodules in the *nfr5* and *symrk* plants (Supplemental Table S2), indicating that IRBG74 nodulation is Nod factor-dependent and requires functional NF receptors for recognition of the Nod factor produced by IRBG74 (Crook et al., 2013; Poinsot et al., 2016) to trigger downstream signal transduction. However, the nodulation performance of the *rinrk1* mutant was different between plants inoculated with IRBG74 and *M. loti*. In *rinrk1* plants inoculated with *M. loti* many white uninfected nodules were formed, with very few pink nodules at the different time points analysed (Fig. 4A and C). In contrast, this hypernodulated-uninfected phenotype was not observed with IRBG74. The *rinrk1* plants infected by IRBG74 developed similar numbers of nodule structures to w. t. plants inoculated with *M. loti* or IRBG74 at 4-6 wpi, with the majority of them pink, indicating effective rhizobial colonization and symbiosis (Fig. 4A and F). The number of pink nodules was significantly higher in plants infected by IRBG74 at 3-6 wpi compared to plants colonized by *M. loti*, indicating that intercellular infection was not impaired in the *rinrk1* mutant. The role of the expolysaccharide receptor *EPR3* is apparently important for both types of infection in *L. japonicus*, since a comparable delayed nodulation phenotype of the *epr3* mutant was observed following inoculation with *M. loti* or IRBG74 (Fig. S2A).

Since the infection process is also controlled by several transcriptional regulators, the role of *Nin, Cyclops, Ern1, Nsp1* and *Nsp2* were tested with IRBG74. The transcription factors *Nin*, *Nsp1* and *Nsp2* are indispensable for the symbiotic process established by IRBG74 and *M. loti*, since the mutants affected in these genes were unable to form nodules (Supplemental Table S2). The nodulation phenotype of *cyclops* plants inoculated with IRBG74 and *M. loti* was similar, and only uninfected white nodules developed (Supplemental Fig. S2B). However, a clear difference was found in the symbiotic performance of the *ern1* mutant. The first pink nodules appeared at 4 wpi with *M. loti* (Fig. 4A and D), while in the presence of IRBG74 these were detected at 3 wpi with fully developed pink nodules at 4 wpi (Fig. 4A, G). This is the reverse of the kinetics shown by w. t. plants wherein pink nodules emerged at 2 and 3 wpi with *M. loti* and IRBG74, respectively (Fig. 1A and B). Additionally, both the total number of nodules and pink nodules was higher in *ern1* mutants with IRBG74 at 3-6 wpi, compared to *M. loti* (Fig. 4A).

**Figure 4.**
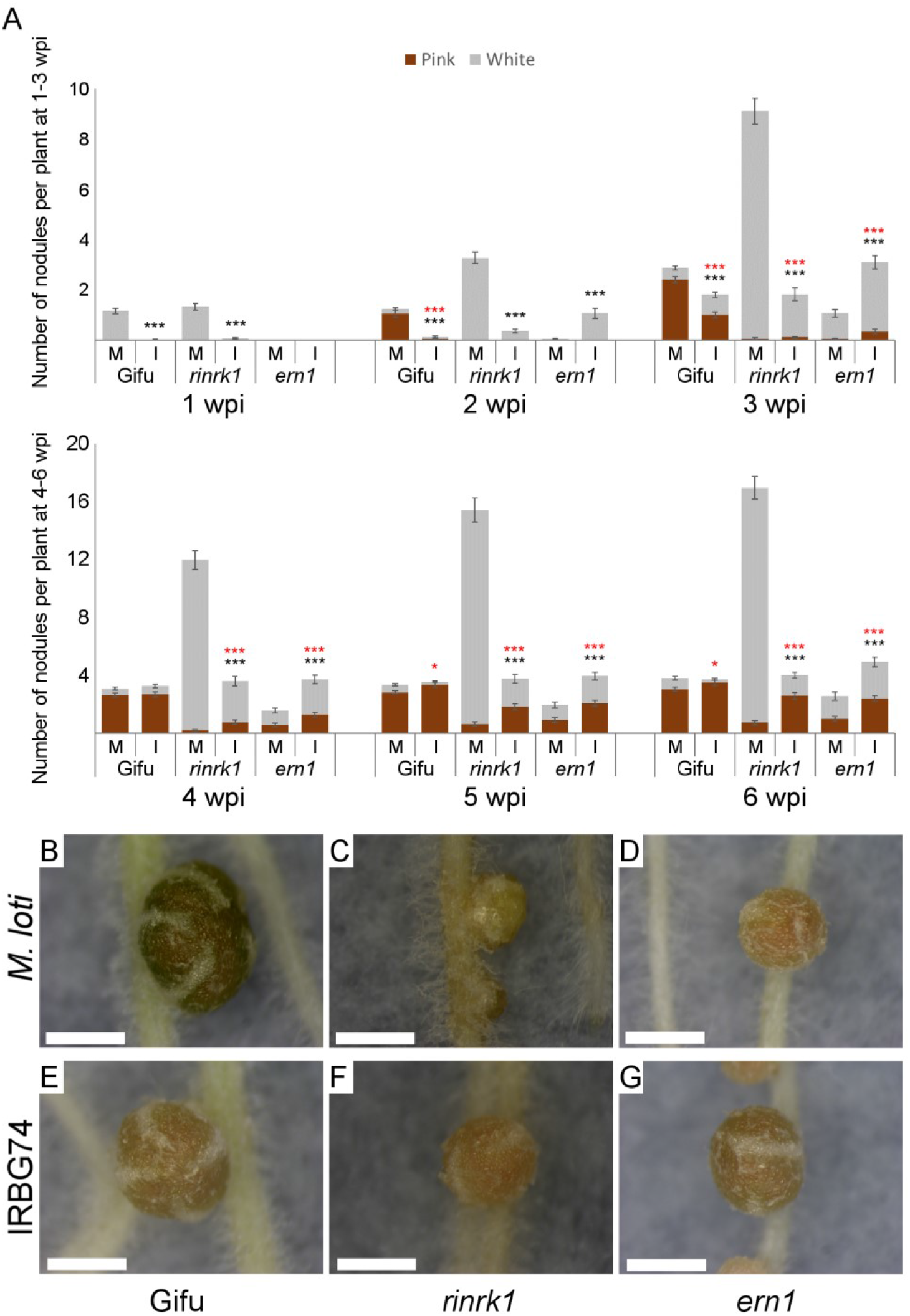
Nodulation phenotype of *rinrk1* and *ern1* mutants. Nodulation kinetics from 1-6 wpi (A) with *M. loti* (M) or IRBG74 (I) in *rinrk1* and *ern1*. Error bars mean SEM. Student’s *t*-test of pink and total number of nodules (red and black asterisks, respectively) between plants inoculated with *M. loti* or IRBG74 in the same genetic background. P-values < 0.05 and 0.001 are marked with one or three asterisks, respectively. n = Gifu 59 (M), 69 (I); *rinrk1* 49 (M), 55 (I); *ern1* 48 (M), 59 (I). Representative images of nodules at 4 wpi with IRBG74 (E, F and G) or *M. loti* (B, C and D) from Gifu (B and E), *rinrk1* (C and F) and *ern1* (D and G). Scale bar 1 cm (G) and 1 mm (H-J).

### Nodulation by IRBG74 is negatively impacted in root hair IT mutants

In *Lotus* several genes participating in a wide variety of molecular processes have been described as important for IT progression. For instance, the *Lotus* mutants affected in the U-box protein *Cerberus* (Yano et al., 2009), the nodule pectate lyase *Npl1* (Xie et al., 2012) or the cytoskeleton component *ScarN* (Qiu et al., 2015), show defects in IT growth and progression. The contribution of these genes to the intercellular infection of IRBG74 was therefore assessed. These mutants were characterized by the formation of a high proportion of white nodules both with *M. loti* and IRBG74 (Supplemental Fig. S2C-E), and few pink nodules were developed in the *npl1* and *scarN* mutants (Supplemental Fig. S2D and E). In all these mutants, the number of nodule structures was reduced when IRBG74 was used as inoculum (Supplemental Fig. S2C-E).

In *Medicago* IT development has been shown to require the function of the *Vapyrin, RPG* and *Cbs* genes. Since the participation of these genes has not been reported in *Lotus*-rhizobia symbiosis, their homolog counterparts were identified in the *Lotus* genome (https://lotus.au.dk/). For *Vapyrin*, two homologous genes named *LjVpy1* (LotjaGi2g1v0091200) and *LjVpy2* (LotjaGi1g1v0646300) (Supplemental Table S1) encoding proteins with 77% and 69% identity to the *Mt*VPY protein, respectively. Similarly, two genes encoding proteins with 61% and 31% amino acid identities to *Mt*RPG were named *Lj*RPG (LotjaGi5g1v0253300) and *Lj*RPG-like (LotjaGi5g1v0086600), respectively (Supplemental Table S1). The *Lj*CBS found (LotjaGi2g1v0126500) showed 78% of protein similarity to *Mt*CBS. Using the LORE1 database, mutant lines with LORE1 insertions were identified and genotyped to obtain homozygous mutant plants. A delay in nodule organogenesis was observed for all the mutants inoculated with *M. loti*, reflected by a reduced number of nodules within the first week after rhizobial inoculation (Supplemental Fig. S2F-J). In response to IRBG74 inoculation, nodulation was also delayed in *vpy1* and *vpy2* (Supplemental Fig. S2G and H), but not in the *rpg* and *rpg-l* mutants, where the number of nodules were similar to the w. t. plants at different time points (Supplemental Fig. S2I and J). Based on the nodule numbers, *vpy1* plants were more severely impacted with IRBG74 (Supplemental Fig. S2G), while the *vpy2* mutant was more affected with *M. loti* (Supplemental Fig S4H). However, at 5 and 6 wpi both the total number of nodules and pink nodules, tended to be similar between the w. t. and mutants. Since *MtVpy* and *MtRPG* have been described as important for IT development in *Medicago*, the number of root hair ITs was recorded in mutants affected in these genes in *Lotus* at 1 wpi with *M. loti*-DsRed. A significant reduction of IT numbers of around 50% in the *vpy1, vpy2* and *rpg* mutants compared to w. t. was observed (Supplemental Table S3).

### Dispensable role of ROS and ethylene in the *Lotus*-IRBG74 symbiosis

It has been described that ethylene plays a positive role in the intercellular infection and nodulation program in the *S. rostrata-Azorhizobium* symbiosis (D’Haeze et al., 2003). To determine the role of this phytohormone in the *Lotus*-IRBG74 symbiotic process, the nodulation kinetics of the double mutant insensitive to ethylene *ein2a ein2b* (Reid et al., 2018) was recorded. The first nodule structures appeared at 2 wpi in the *ein2a ein2b* mutant inoculated with IRBG74, showing a hypernodulation phenotype in the subsequent weeks. However, the total number of nodules induced by IRBG74 was considerably lower than *M. loti* (Supplemental Fig. S2K). In addition, ethylene production was lower in w. t. *Lotus* roots inoculated with IRBG74 in comparison to plants treated with *M. loti* (Supplemental Fig. S3). Taken together, these results show that ethylene is not playing a pivotal role in the infection and organogenesis program triggered by IRBG74 in *Lotus*.

In several legumes it has been shown that reactive oxygen species (ROS) produced by respiratory burst oxidase homolog (RBOH) enzymes are required for intracellular and intercellular infection (Peleg-Grossman et al., 2007; Montiel et al., 2012; Montiel et al., 2016; Arthikala et al., 2017). To address the involvement of these compounds in the symbiotic process induced by IRBG74 in *Lotus*, homozygous mutant lines affected in two *Rboh* isoforms, *LjRbohE* (LotjaGi5g1v0224200) and *LjRbohG* (LotjaGi1g1v0771200), were obtained from the LORE1 database. *LjRbohE* and *LjRbohG* are putative orthologs of *MtRbohA* and *PvRbohB*, previously characterised genes, required for nodule functioning and rhizobial infection in *Medicago* and common bean, respectively (Marino et al., 2011; Montiel et al., 2012). In response to *M. loti* inoculation both *rbohE* and *rbohG* showed a reduced number of nodule primordia and pink nodules at 1 wpi (Supplemental Fig. S2L and M). However, in the ensuing weeks these nodule structures attained similar numbers to w. t. plants at all timepoints tested. The nodulation kinetics of these *rboh* mutants in response to IRBG74 infection was comparable to the w. t. plants (Supplemental Fig. S2L and M), which indicate that these genes are not playing an important role in the intercellular colonization by IRBG74 in *Lotus*.

### Intercellular infection by IRBG74 promotes a distinct transcriptional reprogramming

The nodulation assays of the *Lotus* mutants inoculated with IRBG74 showed that the genetic dependencies for intercellular and intracellular infection modes differ. To further evaluate the signalling pathways involved in intercellular infection, RNAseq transcriptome data were collected and analysed from IRBG74-inoculated *Lotus* roots at 3, 5 and 10 dpi, to cover the infection phase preceding nodule organogenesis. These data were compared to available RNAseq information on *Lotus* roots harvested at 1 and 3 dpi with *M. loti* (Mun et al., 2016; Kamal et al., 2020). In response to *M. loti* and IRBG74 inoculation, a total of 12,637 and 10,947 differentially expressed genes (DEG, P-adjust < 0.5) were identified, respectively (Supplemental Fig. S4A; Supplemental Table S4). The largest transcriptome responses were observed at 1 dpi with *M. loti* (12,534) and at 10 dpi with IRBG74 (9,438) DEG (Supplemental Fig. S4B). A more stringent analysis, with the DEG showing a LOG2FC ≥ 2, revealed that the most important transcriptome response was triggered at 5 dpi IRBG74 (Fig. 5A). Interestingly, only 33% of the DEG by *M. loti* were similarly affected by IRBG74 and a large proportion of the up/down-regulated genes during *M. loti* infection (314: sum of the up and down-regulated) were not similarly affected by IRBG74 (Fig. 5B and C). Additionally, the majority of the genes down-regulated in response to IRBG74 inoculation were not repressed in the *M. loti* transcriptome (Fig. 5B and C).

**Figure 5.**
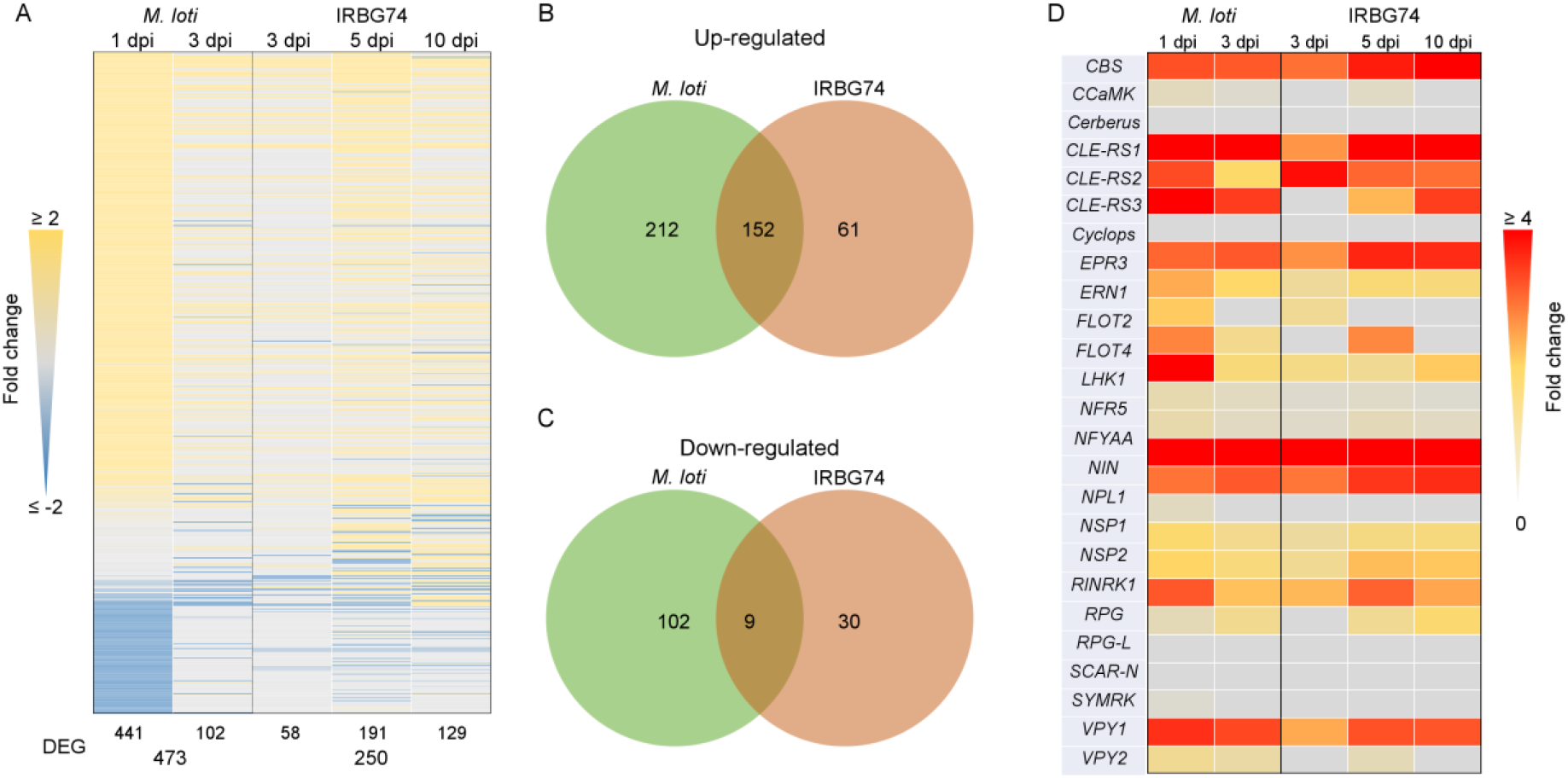
Comparison of the transcriptomic profile of *Lotus* roots during intracellular and intercellular rhizobial infection. A, heat-map expression of DEG (LOG2FC ≥ 2 and a P-adjust < 0.5) in after *M. loti* or IRBG74 inoculation. B and C, Ven diagrams with total number (*M. loti*: 1 and 3dpi; IRBG74: 3, 5 and 10 dpi) of the up/down regulated DEG during intracellular (*M. loti*) and intercellular (IRBG74) colonization. D, heat-map expression of known SYM genes significantly (P-adjust < 0.5) induced after rhizobial perception.

The nodulation kinetics of *Lotus* inoculated with IRBG74 revealed that nodule organogenesis was delayed with respect to plants inoculated with *M. loti*. To determine if this delay was linked to a deficient induction of the early symbiotic signalling pathway, the expression profile of several genes known to be involved in infection and/or nodule organogenesis were analysed using the transcriptome data set. Most of the known symbiotic genes tested were induced by *M. loti* or IRBG74 (Fig. 5D), but when the same time point was compared (3 dpi), several genes were less upregulated in response to IRBG74 inoculation (Fig. 5D).

### Cytokinin signalling is differentially regulated in response to IRBG74

This work showed that *Ern1*, a transcription factor implicated in cytokinin signalling (Cerri et al., 2017; Kawaharada et al., 2017), has a less relevant role during intercellular infection by IRBG74. These results suggest that this mode of infection triggers a different transcriptional response of cytokinin-related genes compared to intracellular infection. To validate this hypothesis, a transcriptome heat-map was created for genes involved in cytokinin synthesis and regulation and significantly affected by *M. loti* or IRBG74 inoculation (Fig. 6). This approach confirmed the different gene-expression response of several components of cytokinin regulation during IRBG74 infection. Particularly, genes encoding the cytokinin degrading enzymes *Ckx3, Ckx9*, the response regulator involved in cytokinin signalling *RR11a* and the cytokinin biosynthesis gene *Ipt2* were poorly or insignificantly induced during intercellular infection, relative to their evident up-regulation after *M. loti* inoculation (Fig. 6). *M. loti* triggered a significant reduction in the expression levels of *Ckx2, Ckx8, Lhk2, RR3a, RR19*, and *Log*. However, most of these genes were not significantly down-regulated by IRBG74 colonization. By contrast, the cytokinin biosynthesis gene *Cyp735a*, which converts iP to tZ type cytokinins, showed a strong up-regulation after IRBG74 colonization at 5 and 10 dpi, whereas its gene expression was only slightly altered during intracellular infection at early timepoints (Fig. 6).

**Figure 6.**
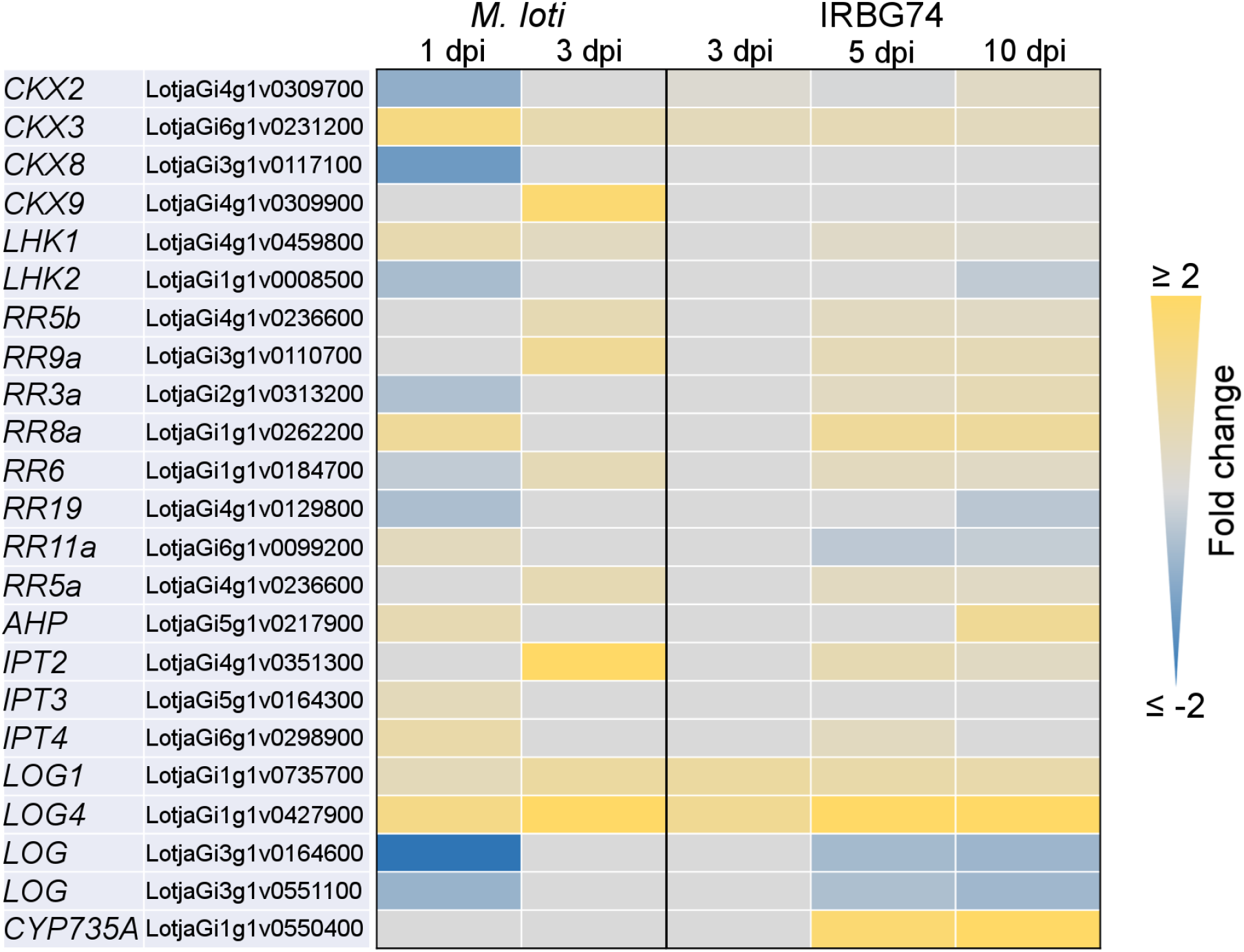
Heat-map of DEG encoding cytokinin related proteins in *Lotus* roots after rhizobial inoculation. The heatmap highlights the differences in the expression profile of DEG (log2FC ≥ 0.5, P adjust < 0.5) involved in the metabolism, perception, signalling and synthesis of cytokinins in response to *M. loti* or IRBG74 infection.

### *Cyp735a, Ipt4* and *Lhk1* are relevant players in the *Lotus*-IRBG74 symbiosis

In the *Lotus-M. loti* symbiosis, the cytokinin receptor mutant *lhk1* shows a delayed and reduced nodulation (Murray et al., 2007), while the *ipt4* and *cyp735a* mutants show minor or insignificant phenotypes, respectively (Reid et al., 2017; Supplemental Fig. S5). The RNAseq data presented in this study indicates that these genes are differentially regulated by *M. loti* or IRBG74. In order to determine the relevance of these cytokinin-related genes in the *Lotus*-IRBG74 symbiosis, nodulation kinetics at 1-6 wpi were scored for the *cyp735a, ipt4* and *lhk1* mutants after IRBG74 inoculation. The nodulation capacity of both *cyp735a* and *ipt4* mutants was substantially reduced, although with different symbiotic phenotypes. The *cyp735a* mutant developed similar numbers of nodules to w. t. plants, at 2 and 3 wpi (Fig. 7A), but at 4-6 wpi *cyp735a* formed more nodule-like structures, most of them uninfected white nodules (Fig 7C, right panel). By contrast, at 2-4 wpi the total number of nodules in *ipt4* was lower compared to w. t. plants (Fig. 7A). The number of pink nodules was reduced at all time points tested and these comprised a mixture of pink and pale pink nodules (Fig. 7A and D, right panel). Interestingly the *lhk1* mutant was unable to develop any nodule structure in the presence of IRBG74 (Fig. 7A). This drastic symbiotic phenotype contrasts with the *lhk1* plants inoculated with *M. loti*, where several pink nodules were observed at 6 wpi (Supplemental Fig. S6). These results further demonstrate the different cytokinin regulation during intercellular infection in *Lotus*, whereby *Cyp735a*, *Ipt4* and *Lhk1* are important players for this type of rhizobial infection (Fig. 8).

**Figure 7.**
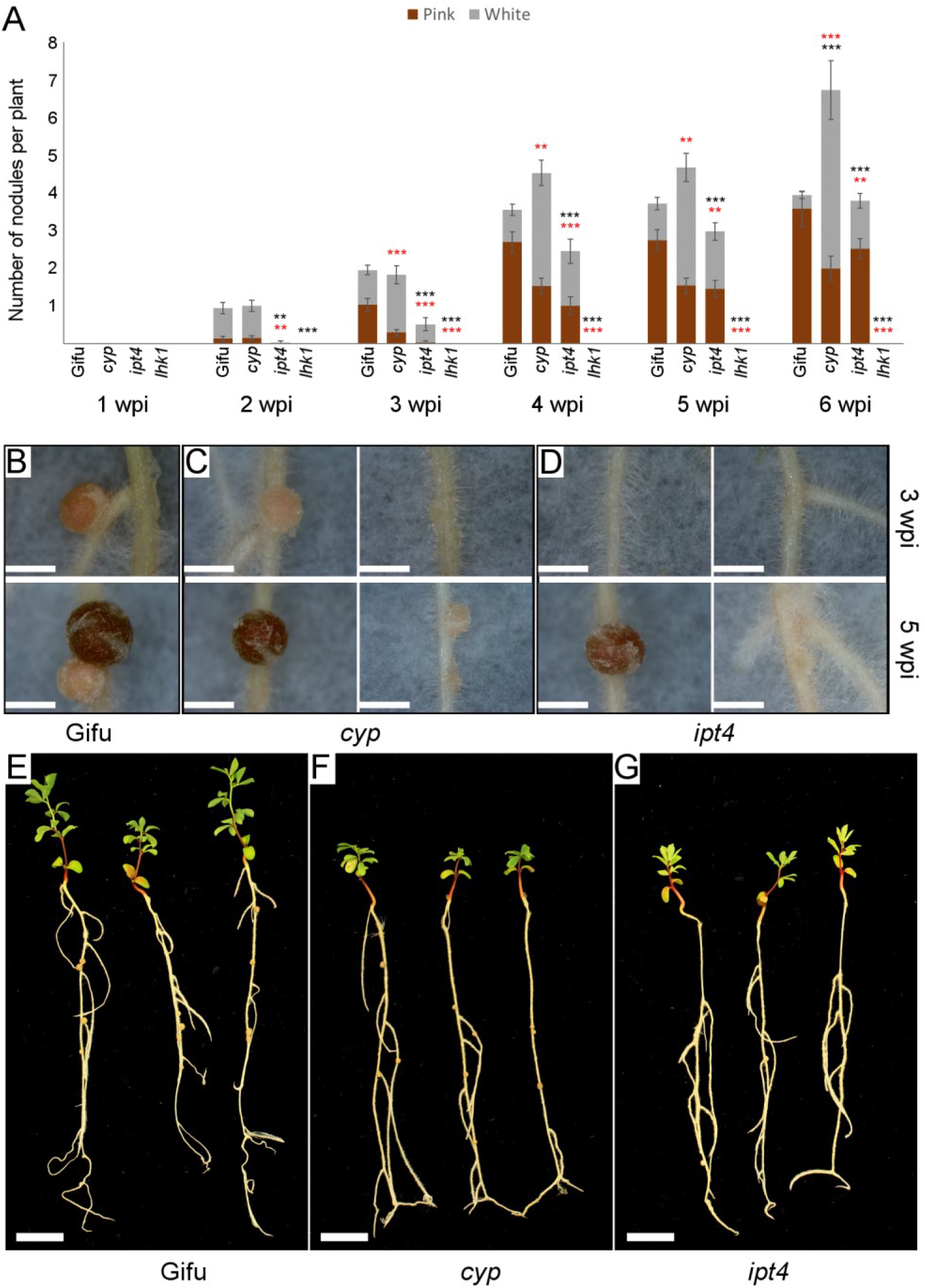
Symbiotic phenotype of *Lotus* mutants affected in cytokinin-related genes. A, nodulation kinetics of *cyp735a, ipt4* and *lhk1* plants from 1 to 6 wpi with IRBG74. Error bars mean SEM. Student’s *t*-test analyses of pink nodules and total number of nodules (red and black asterisks, respectively) between w. t. and mutant plants inoculated with IRBG74. P-values < 0.01 and 0.001 are marked with two or three asterisks, respectively. n = 49 (Gifu), 86 (*cyp735a*), 53 (*ipt4*), 55 (*lhk1*). Phenotype of nodules developed in w. t. (B), *cyp735a* (C) and *ipt4* (D) plants at 3 (upper panel) and 5 wpi (lower panel) with IRBG74. Representative images of w. t. (E), *cyp735a* (F) and *ipt4* (G) plants at 6 wpi with IRBG74. Scale bar, 1 mm (B-D) and 1 cm (E-G).

**Figure 8.**
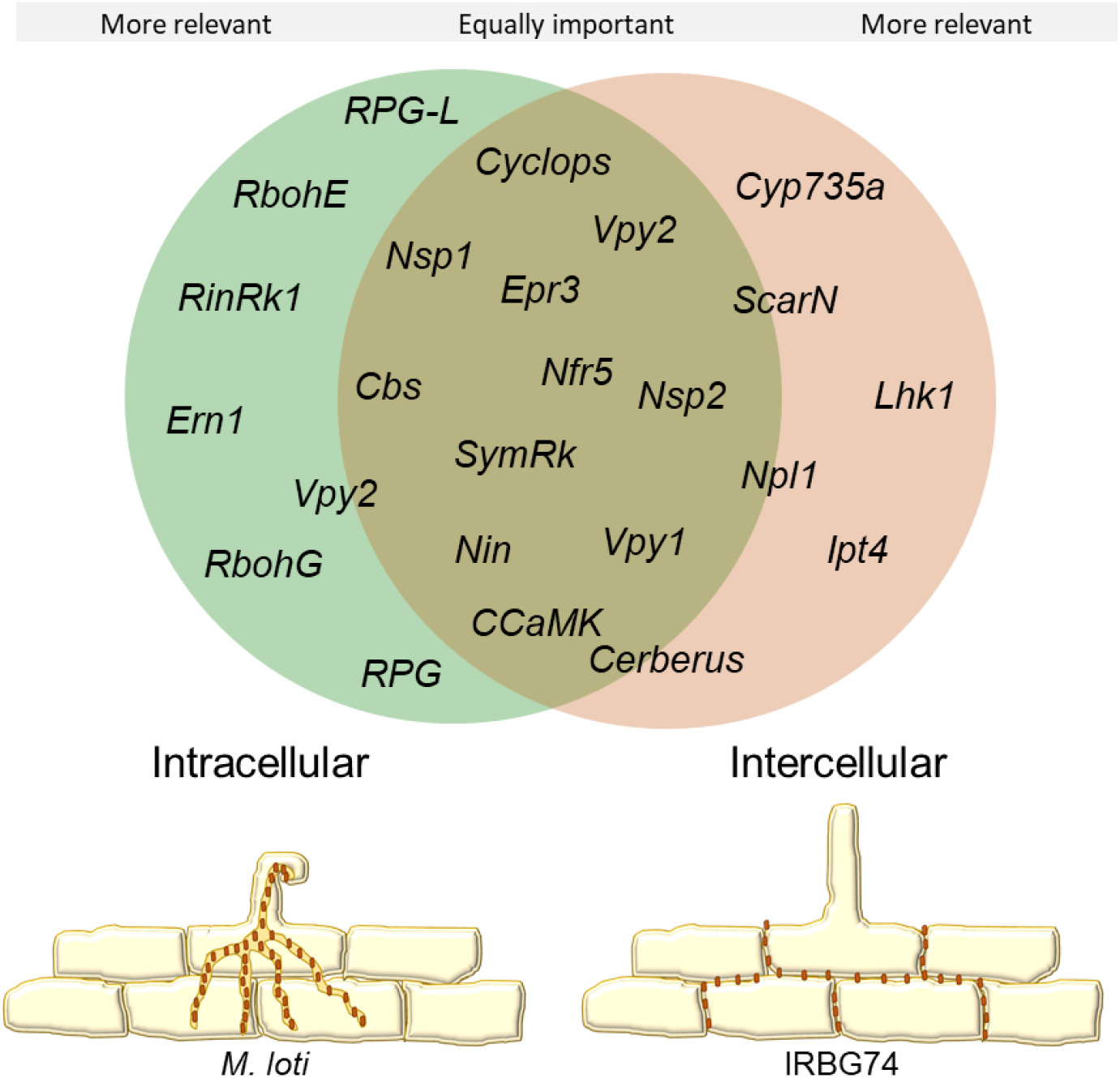
Comparative model of gene requirements in the intracellular and intercellular symbiotic program in *Lotus*. *Lotus* recruits a core of essential genes for the root nodule symbiosis, regardless the type of infection mechanism employed by rhizobia. Nonetheless, certain players are particularly more relevant depending on the mode of colonisation. Intercellular infection in *Lotus* appears to be more sensitive to the absence of certain cytokinin-related genes.

## Discussion

### *Lotus*-IRBG74 symbiosis: a novel working model to study intercellular infection

Intercellular colonization of legumes by rhizobia occurs via a variety of entry modes. Bacteria can penetrate through middle lamellae of root hairs, cracks at emergent lateral roots or between epidermal cells (Subba-Rao et al., 1995; Gonzalez-Sama et al., 2004; Goormachtig et al., 2004; Bonaldi et al., 2011). Diverse *Lotus* spp. exhibit this infection mechanism, which leads to the formation of either ineffective or nitrogen-fixing nodules, depending on the growth conditions and rhizobial partner (Ranga Rao, 1977; James and Sprent, 1999; Liang et al., 2019). Previously, it was described that under certain mutant background conditions and with a low frequency, *Lotus* roots can be intercellularly colonized by *M. loti* (Madsen et al., 2010) by a NF-independent mechanism. Here we show that IRBG74 massively accumulates on the root surface of *Lotus* roots and invades the roots between the epidermal and root hair cells. This intercellular invasion transforms into transcellular infection threads in inner root cell layers and the nodule. A similar scenario has been described in *S. rostrata*, where the infection pocket, formed by an intercellular invasion of certain *Azorhizobium* spp., is followed by transcellular infection threads (Goormachtig et al., 2004). Although IRBG74 promoted nodule formation in *Lotus*, the organogenesis and infection programs were delayed, compared to the symbiotic proficiency of *M. loti*. This delay is apparently not related to the infection mode, since the infection and nodule organogenesis program in *A. hypogaea*, which is also intercellularly invaded, exhibit comparable kinetics to those observed in the model legumes *Lotus* or *Medigaco*, wherein root hair ITs are formed.

### Common gene dependencies of intercellular and intracellular colonization

The characterisation of several legume mutants has identified the molecular players of the symbiotic pathway, from the early signalling to nodule organogenesis. NF receptors are indispensable for initiating symbiotic signalling, since *Lotus* mutants disrupted in these genes do not show any symbiotic response (Madsen et al., 2003; Radutoiu et al., 2003). The nodulation test of *Ljnfr5* with IRBG74 revealed that functional NF receptors are required for the *Lotus*-IRBG74 symbiosis. Likewise, both intercellular rhizobial infection and nodule organogenesis is a NF-dependent process in the *S. rostrata-A. caulinodans* relationship (Capoen et al., 2010), but there are also intercellular processes whereby a NF-independent mechanism can lead to nitrogen-fixing nodules in certain legumes (Giraud et al., 2007; Madsen et al., 2010; Ibanez and Fabra, 2011). This study revealed a genetic machinery that is equally important for both types of infection modes, since a similar detrimental impact in the nodulation process was observed in the *nfr5, symrk, ccamk, cyclops, nin, nsp1, nsp2 epr3, cbs* and *vpy1* mutants whether *M. loti* or IRBG74 were used as inoculum in *Lotus* (Fig. 8). However, the intracellular symbiotic program was more affected in the *rinrk1, ern1, rbohE, rbohG, rpg, rpg-like*, and *vpy2* mutants. One interpretation is that these genes are more important for formation of root hair ITs than transcellular ITs. Previously, it was described in *A. evenia* that SYMRK, CCaMK and the histidine kinase HK1 are required both in the intracellular and intercellular infection by *Bradyrhizobium* (Fabre et al., 2015), reinforcing the notion of a common genetic repertoire for these types of rhizobial infection. Likewise, SYMRK and CCaMK are required for intercellular infection and nodulation in the IRBG74-*Lotus* symbiosis. However, the symbiotic process through intercellular colonization is apparently more sensitive to the absence of certain genes. *cerberus*, *npl1* and *scarN* mutants developed numerous white nodules, frequently uninfected after *M. loti* inoculation (Yano et al., 2009; Xie et al., 2012; Qiu et al., 2015), but when IRBG74 was used as inoculum, nodule development and the number of pink nodules were severely reduced in these mutants. This suggests that actin rearrangement play an important role in formation of cortical and transcellular infection threads and that initiation of ITs from intercellular infection pockets is more dependent on actin rearrangement.

As mentioned above, there are few reports describing the molecular components required for intercellular infection. One of them, revealed the positive role of ROS to induce cell death during the crack entry infection of *Azorhizobium* in *S*. *rostrata*. Deprivation of ROS production by applying diphenyleneiodonium chloride, an inhibitor of the ROS-producing enzymes RBOHs (for respiratory burst oxidase homolog), prevents rhizobial colonization in this legume (D’Haeze et al., 2003). Similarly, these genes have been implicated in the IT development during intracellular rhizobial infection in *Medicago* and *P. vulgaris* (Peleg-Grossman et al., 2007; Montiel et al., 2012; Arthikala et al., 2017). Likewise, the nodulation program was delayed in *Lotus* mutants disrupted in *RbohE* or *RbohG* genes after inoculation with *M. loti*, however, the nodulation kinetics in these mutants was unchanged by IRBG74. Inhibition of ethylene synthesis or perception has a negative effect in the nodulation and intercellular infection induced by *Azorhizobium* in *S. rostrata* (D’Haeze et al., 2003). In contrast, ethylene plays a negative role in the *Lotus*-*M. loti* symbiosis. The *Ljein2a Ljein2b* double mutant that exhibits complete ethylene insensitivity is hyperinfected and hypernodulated by *M. loti* (Reid et al., 2018). Unlike, the intercellular symbiotic process in *S. rostrata*, were ethylene is required for nodulation, the *Ljein2a Ljein2b* mutant was hypernodulated by IRBG74, indicating that this phytohorme is not essential for the IRBG74 intercellular infection.

### Distinctive cytokinin signalling program during intercellular infection

The dual role of cytokinins, as positive and negative regulators of nodule development and rhizobial infection, respectively, makes them key phytohormones in the legume-rhizobia symbiosis (Miri et al., 2016). The *lhk1* mutant belatedly develops a reduced number of pink nodules in response to *M. loti* infection, but when IRBG74 is used as inoculum the mutant is unable to form nodules up to 6 wpi. In the intracellular infection mediated by *M. loti*, it has been suggested that other cytokinin receptors are sufficient to induce nodule organogenesis in the absence of *Lhk1* (Murray et al., 2007). However, the signalling pathway triggered by LHK1 is indispensable in the *Lotus*-IRBG74 symbiosis. Conversely, the *ern1* mutant displayed improved symbiotic performance with IRBG74, which further confirms that depending on the type of rhizobial infection program, distinct signalling pathways are triggered. This was reflected in the different transcriptomic responses of genes involved in the synthesis, perception signalling and metabolism of cytokinin induced by *M. loti* or IRBG74. Similarly, the *A. hypogaea* roots intercellularly infected by rhizobia, exhibit a distinctive transcriptome profile of genes involved in cytokinin metabolism and signalling, when compared to the expression pattern of orthologue genes in legumes intracellularly colonized (Peng et al., 2017; Karmakar et al., 2019). It has been shown that different cytokinin biosynthesis genes are induced during intracellular colonization of *M. loti* in *Lotus* roots (Reid et al., 2016; Reid et al., 2017). Although the cytokinin *trans*-hydroxylase *Cyp735a* is highly induced by *M. loti*, the nodulation performance is not significantly affected in *Lotus* plants disrupted in this gene (Reid et al., 2017). By contrast, nodule organogenesis is delayed and reduced in *cyp735a* mutants inoculated with IRBG74. *Cyp735a* encodes cytochrome P450 monooxygenases (P450s) that catalyze the biosynthesis of trans-Zeatin, an isoprenoid cytokinin compound (Takei et al., 2004), indicating that tZ cytokinins play a more relevant role during intercellular colonization. The mutant affected in the isopentenyl transferase 4 (*Ipt4*) gene, showed a mild impact in the nodulation capacity with *M. loti* (Reid et al., 2017). However, in response to IRBG74 inoculation, the development of nodules was evidently delayed, and the number of pink nodules reduced. IPT is placed in the first step during isoprenoid cytokinin biosynthesis, giving rise to iP riboside 50-diphosphate (iPRDP) or iP riboside 50 - triphosphate (iPRTP) intermediates, which can be converted by CYP735a to tZ cytokinins (Hwang and Sakakibara, 2006). The delayed nodulation and enhanced phenotypes of the cytokinin related mutants may indicate that the peak of cytokinin triggered by the intercellular programme is lower or more disperse in the root relative to the highly localised cytokinin signalling achieved in the intracellular infection modes.

## Conclusions

The intracellular colonization has been extensively described at the cellular level in recent decades, but there is little knowledge about the molecular players controlling the intercellular invasion. Different approaches revealed that the cytokinin signalling pathway is apparently a key difference to be further analysed. Similarly, other components seem to be differentially relevant for this type of infection. For instance, the *rinrk1* receptor mutant showed a better nodulation performance with IRBG74 compared to *M. loti*. This latter indicates that during intercellular infection, certain uncharacterized ligands are not entirely necessary for rhizobial colonization.

## Material and Methods

### Germination and Nodulation assays

*Lotus* seeds of accession Gifu (Handberg and Stougaard, 1992) were scarified with sandpaper were surface sterilized with 0.3% of sodium hypochlorite for 10 minutes and then washed 5 times with autoclaved distilled water, to remove traces of chlorine. The washed seeds were incubated overnight at 4 °C and then transferred to square petri dishes for germination at 21 C. For monitoring nodulation kinetics, three days post-germination seedlings (n ≥ 20 plants per condition) were placed into petri dishes with 1.4% B&D agar slant covered with filter paper and inoculated with the respective rhizobial strains (OD_600_ = 0.05). After rhizobial inoculation the number of white (bumps and nodule primordia) and pink nodules were recorded weekly until 6 weeks post inoculation (wpi) with a stereomicroscope. The plants were harvested at 6 wpi to measure the fresh weight and length of the aerial part. LORE1 lines (Urbanski et al., 2012; Malolepszy et al., 2016) disrupted in the genes of interest were ordered from the LORE1 database (https://lotus.au.dk/) and genotyped with allele-specific primers to obtain homozygous mutants following the database instructions (Mun et al., 2016). The gene IDs and the corresponding LORE1 IDs are shown in Supplemental Table S1.

### Infection phenotyping using confocal microscopy

Three days old seedlings of *Lotus* (accession Gifu) (Handberg and Stougaard, 1992) were placed on ¼ B&D plates and inoculated with fluorescently labelled *M. loti* R7A (Kelly et al., 2013) or IRBG74 strains (Cummings et al., 2009), obtained through transformation with the constitutive DsRED expressing plasmid pSKDSRED (Kelly et al., 2013). The roots were harvested from the plates at different time points. To enable observations in deeper part of the tissue, fluorescent compatible clearing protocol was used as described before (Nadzieja et al., 2019). Cleared roots were visualized by confocal microscopy with the following excitation lasers/emission cutoffs: 405/408–498 nm (autofluorescence), 561/517–635 nm (DsRed). For 3D projections, Fiji ImageJ (Schindelin et al., 2012) was used to create animation frames, which then were rearranged using Adobe Photoshop CC into final projections.

### Nodule histology analysis

Young and mature nodules at 3 wpi with IRBG74 or *M. loti* were detached from *Lotus* roots, sliced in half, and incubated overnight in fixative solution (2.5% glutaraldehyde, 0.1 M sodium cacodylate pH 7). The fixed nodules slices were embedded in acrylic resin and sectioned for light microscopy (James and Sprent, 1999; Madsen et al., 2010).

### RNAseq of *Lotus* roots and bioinformatics

The susceptible infection zone of *L. japonicus* roots by IRBG74 (elongation and maturation zone of the root) was cut and freeze in liquid nitrogen from seedlings at 3, 5 and 10 dpi with IRBG74 (O. D. 600 = .0.05) or mock-treated (water) at the same time points. Total RNA was isolated and DNA contamination was removed by DNAse treatment. Library preparations using randomly fragmented mRNA was performed by IMGM laboratories (Martinsried, Germany) and sequenced in paired-end 150 bp mode on a Illumina NovaSeq 6000 instrument.

A decoy-aware index was built for Gifu transcripts using default Salmon parameters and reads were quantified using the --validateMappings flag (Salmon version 0.14.1; Patro et al., 2017). Normalised expression levels and differential expression testing was performed using the R-package DESeq2 version 1.20 (Love et al., 2014) after summarising gene level abundance using the R-package tximport (version 1.8.0).

### Accession numbers

The RNAseq reads associated with this study are available in the SRA under bioproject accession number PRJNA632725. The *L. japonicus* (accession Gifu and MG20) gene identifiers are shown in Supplemental Table S1.

### Large datasets

The calculated expression values and statistics of the RNAseq data are included as Supplemental Table S4.

## Acknowledgements

We thank the assistance in the greenhouse of Finn Pedersen and Nanna Walther. We thank Dr.Fang Xie for providing *rinrk1* seeds.

## Supplemental Material

Supplemental Figure S1. Nodule cell occupancy in *Lotus* nodules colonized by *M. loti* or IRBG74.

Supplemental Figure S2. Nodulation performance of *Lotus* mutants.

Supplemental Figure S3. Ethylene production in *Lotus* roots inoculated with rhizobia.

Supplemental Figure S4. DEG in *Lotus* roots after rhizobial inoculation.

Supplemental Figure S5. Phenotype of *cyp735a* and *ipt4* mutants inoculated with *M. loti*.

Supplemental Figure S6. Phenotype of *lhk1* plants inoculated with *M. loti* or IRBG74.

Supplemental Table S1. List of *Lotus* mutants used in this study.

Supplemental Table S2. *Lotus* mutants with a Nod-phenotype in response to *M. loti* or IRBG74 inoculation.

Supplemental Table S3. Number of ITs per plant at 1 wpi with *M. loti*.

Supplemental Table S4. RNAseq expression data of *Lotus* roots inoculated with *M. loti* (1 and 3 dpi) or IRBG74 (3, 5 and 10 dpi).

Supplemental Video S1. 3D-projection of *Lotus* roots infected by IRBG74 at 1 and 2 wpi.

